# Beyond thresholds: a fully Bayesian framework for quantifying allele count evidence for variant pathogenicity

**DOI:** 10.64898/2026.02.09.704882

**Authors:** Fedor Konovalov

**Affiliations:** Independent Clinical Bioinformatics Laboratory, Moscow, Russia

## Abstract

Allele count data from affected individuals and population controls are central to variant interpretation, yet their evidential meaning is often mediated by discrete thresholds and implicit assumptions. This work introduces a fully quantitative Bayesian framework for dominant rare disease genetics in which all allele count evidence is summarized by a single quantity, the Bayes factor, that evaluates the probability of observing the same data under two explicitly defined competing models. Rather than replacing individual ACMG/AMP pathogenicity criteria, the Bayes factor provides a unified measure that naturally incorporates evidence in both the pathogenic and benign directions. The framework accounts for variation in affected cohort size, penetrance, disease prevalence, and assay error rates, allowing these biologically and technically meaningful quantities to be specified directly instead of absorbed into fixed cutoffs. Application to a non-Finnish European population shows that the dependence of the Bayes factor on observed allele counts is strongly shaped by how the affected cohort is defined and by false positive rates in control datasets. Across representative scenarios, Bayes factor values are broadly compatible with established allele count criteria combinations expressed on odds-ratio scales under typical parameterizations, while remaining tunable beyond these defaults.

## 1 Introduction

Variant interpretation in medical genetics is the process of translating diverse lines of evidence into a judgment about whether a genetic variant is pathogenic.

Historically, this process was largely a form of art: decisions were made within individual laboratories, informed by accumulated experience with specific genes and detailed knowledge of the epidemiology and clinical presentation of particular genetic disorders (Giles et al. 2021). The widespread adoption of next-generation sequencing (NGS) fundamentally altered this landscape by producing large volumes of novel genetic variation across the genome, far beyond gene- or disease-focused scopes (Koboldt et al. 2013; Yang et al. 2013; Turro et al. 2020). In response, a community effort was undertaken to formalize variant interpretation into a unified set of criteria, culminating in the widely adopted 2015 ACMG/AMP guidelines (Richards et al. 2015). These guidelines were specifically described as consensus- and opinion-based, rather than derived from a formal underlying theory (Richards et al. 2015).

Since their introduction, the criteria have evolved along three main axes: (i) reformulation into alternative but largely equivalent guideline systems (Matthijs et al. 2016; Nykamp et al. 2017; Houge et al. 2022; Durkie et al. 2024); (ii) refinement of individual criteria through ClinGen expert guidance (Rehm et al. 2015; Clinical Genome Resource 2026a); and (iii) expansion into gene- or disease-specific rule sets developed by Variant Curation Expert Panels (VCEPs) (Clinical Genome Resource 2026b), in some cases highly detailed (Fortuno et al. 2025).

In parallel, there has been a sustained effort to move variant interpretation toward a more quantitative and scientifically grounded foundation. An important step in this direction was the recognition that the ACMG framework can be interpreted as an implicit Bayesian system for weighing evidence for and against pathogenicity (Tavtigian, Greenblatt, et al. 2018). This insight enabled the mapping of existing criteria onto a calibrated odds-based point system without redefining the criteria themselves (Tavtigian, Harrison, et al. 2020). However, both the thresholds and the relative weights of individual criteria remain largely community-derived. It has long been acknowledged that these parameters are context-dependent and may require adjustment across diseases, genes, or clinical ascertainment scenarios.

For some evidence types, formal statistical modeling has already been introduced. For example, modern aggregate missense pathogenicity scores have been argued to carry more information than is currently accommodated by generic criteria, opening new opportunities about their appropriate calibration and future role (Pejaver et al. 2022; Bergquist et al. 2025). Related efforts have also systematized functional and *de novo* evidence in quantitative or likelihood-ratio terms (Brnich et al. 2020; Samocha et al.2014).

Allele counts observed in affected individuals and in control populations constitute one of the most direct forms of evidence in variant interpretation, as they reflect joint observations of genotype and phenotype in humans. Within the ACMG/AMP framework, such evidence is encoded in several criteria: PS4 and PM2 support pathogenicity, whereas BA1, BS1, and in limited circumstances BS2 support benignity. Importantly, BS2 requires well-documented observations in healthy adults and is therefore not generally applicable to large, population-based control datasets.

The impact of allele-count-based criteria on pathogenicity assessment spans a wide range. Even in the original 2015 guidelines, PS4 was accompanied by caveats, optional odds-ratio-based testing, and provisions for modifying its strength. These ideas were subsequently formalized in quantitative implementations such as the CardioDB PS4 calculator (Bowman 2026). An alternative approach was implemented in the PS4-LRcalc application, which expresses PS4 evidence as a likelihood ratio and can output both categorical and Tavtigian-scaled quantitative strengths (Rowlands et al. 2024). On the benign side, BS1 has been substantially refined to allow variable frequency thresholds that depend on control sample size, penetrance, and allelic or genetic heterogeneity (Whiffin et al. 2017).

Taken together, these developments motivate the synthesis of allele count evidence into a unified framework. Such a framework should satisfy three key requirements. First, it should be inherently quantitative: fixed thresholds inevitably compress information and may become increasingly miscalibrated as sample sizes grow. Second, it should be flexible: allele-frequency-based criteria are among the most frequently modified by VCEPs, reflecting genuine biological diversity across genes and diseases, and a suitable model must therefore expose biologically meaningful, tunable parameters. Third, it should be firmly grounded in population genetics, with assumptions stated rather than implicit.

In this study, this methodology is developed for autosomal dominant variants in the context of rare genetic disease.

## 2 Theoretical approach

In the Bayesian view, the same observed allele counts are informative under both pathogenic and neutral hypotheses, and their significance is encoded in how well each hypothesis is supported by the data. This naturally suggests evaluating allele count evidence in terms of how strongly the observed data favor a pathogenic explanation over a neutral one, while accounting for uncertainty in population- and variant-specific parameters.

This comparison is formalized by the Bayes factor, which quantifies the relative support that the data provide for two competing models. The Bayes factor is defined as the ratio of the marginal likelihoods of the two models being compared, with unknown parameter values integrated over their admissible ranges. Let ℳ_*p*_ denote a model under which the variant is pathogenic and ℳ_*n*_ a model assuming full neutrality. For the empirical data 𝒟, the Bayes factor is:

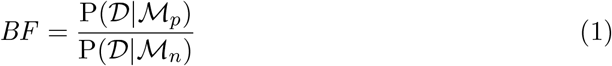

### 2.1 Marginal likelihood under the pathogenic model

The marginal likelihood of a model is the total probability of observing the given data under the model, obtained by integrating the likelihood function P(𝒟| ℳ) over the space of unknown model parameters *θ* weighted by their probability density. For the pathogenic model ℳ_*p*_ applied to autosomal dominant variants:

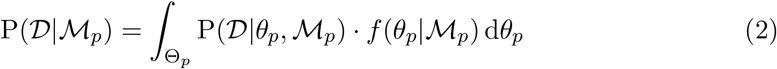

Here, the integration domain Θ_*p*_ is defined as a rectangular region ℛ in parameter space, with its axes corresponding to the true (unknown) total population allele frequency of the variant *x* and the mutation rate *µ*:

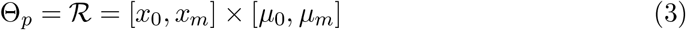

The probability of observing the data, i.e. allele counts, is modeled as the product of two binomial probabilities of detecting *k* variant alleles in *n* chromosomes with rate *π* in the affected sample (*d*) and the control sample (*c*), respectively, assuming their independence:

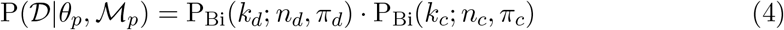

In the absence of detection errors, the binomial rates *π* would coincide with the true allele frequencies in the corresponding cohorts. In practice, however, depending on type and location of the variant, sequencing artifacts may contribute materially to observed allele counts, particularly in large datasets. To account for both type I and type II errors, the binomial rates *π* are derived from the true cohort-specific allele frequencies *x*^*′*^ by introducing sensitivity *Se* and specificity *Sp*, which may differ between affected and control samples:

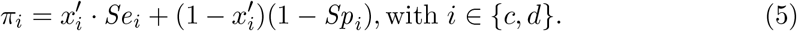

The cohort-specific allele frequencies 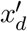 and 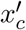 are functions of a single underlying value, the total population allele frequency *x*. This relation depends on the disease prevalence *R*, the penetrance *K* (defined as the fraction of carriers who are affected), and an additional parameter, the inclusion ratio *I* ∈ [0, 1]. The inclusion ratio *I* denotes the fraction of truly affected individuals who are nonetheless present in the control cohort, reflecting incomplete diagnosis or deliberate inclusion of mild phenotypes during control enrollment. Together, these parameters determine the expected allele frequencies in the cohorts as follows:

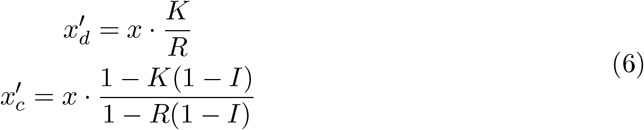

A deterministic estimate of the total population frequency of a fully dominant deleterious variant is given by *x* = *µ/h* (Crow and Kimura 1970), where *µ* denotes the mutation rate and *h*, here, denotes the heterozygote selection coefficient. Modeling based on Exome Aggregation Consortium (ExAC) data has shown that, under moderate-to-strong selection, the effects of human demography and genetic drift introduce only limited deviations from this estimate, becoming pronounced primarily at low values of *h* (Weghorn et al. 2019). A classic extension of the deterministic model to finite populations was provided by Nei 1968, who showed that under strong selection allele frequencies can be approximated by a gamma distribution that preserves the same mean *µ/h* while introducing variance inversely proportional to the effective population size *N* . This distribution is parameterized by shape *α* and rate *λ*, which depend on the mutation rate *µ*, effective population size *N*, and fitness reduction in heterozygotes *h* (with the homozygous state assumed to be lethal). Combined with a probability density for the mutation rate, discussed below, the resulting joint distribution is given by:

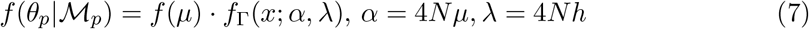

Since analytical evaluation of Eq. (2) is not feasible, numerical methods are used to integrate over *µ* and *x*. The gamma distribution probability density function (PDF), however, poses a challenge for numerical integration when *α* is small, owing to a singularity at *x* → 0. Special care is therefore required to ensure that integration errors remain controlled. This singularity can be removed by a straightforward application of basic integration rules: subtracting from the integrand the same joint PDF multiplied by the binomial probabilities evaluated at *x* = 0, and then adding this term back as a separate integral, ensures that the effective power of *x* in the first integrand is strictly positive. For notational convenience, the binomial probabilities of observing the allele counts in the affected and controls are hereafter denoted by 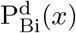 and 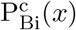, respectively, as being dependent on the population allele frequency *x* via error-adjusted rates *π*_*d*_ and *π*_*c*_; *ϕ* denotes the set of fixed gamma parameters other than *µ*:

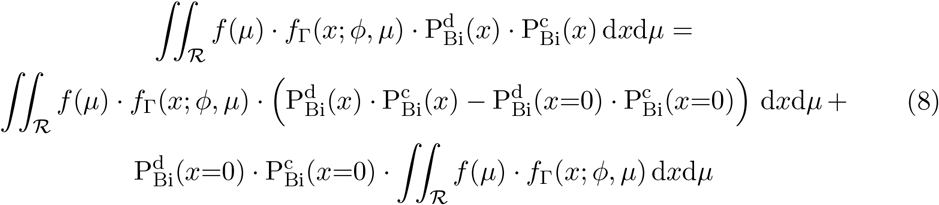

The first integral can then be calculated safely using numerical cubature over ℛ. At

*x* = 0, the integrand may evaluate to an undefined value for some parameter combinations because the gamma PDF is still computed internally; however, since the integrand as a whole approaches zero as *x* → 0, its value can be set to zero at that point without affecting the result. The second term can be evaluated iteratively, with the inner integral corresponding to the cumulative distribution function (CDF) of the gamma distribution, thereby reducing the remaining integration to a single dimension:

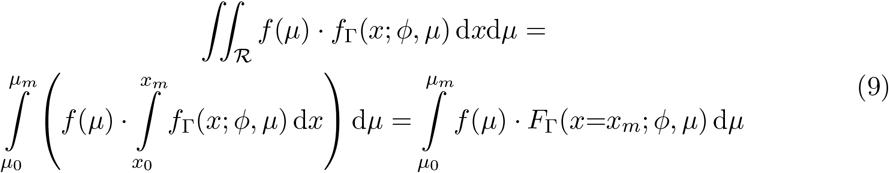

### 2.2 Marginal likelihood under the neutral model

Under neutrality, simple closed-form expressions are generally inadequate for human populations since they are far from equilibrium. The action of genetic drift depends strongly on historical population growth rates and bottlenecks, forming a complex distribution of allele frequencies (site frequency spectrum, SFS) when combined with natural variation in mutation rates. However, if the SFS of a hypothetical reference population of size *N* is known, then it is straightforward to derive likelihoods for observed allele counts in affected and control cohorts sampled from that population using hypergeometric formulas, or, more practically, their binomial approximations. For evaluating marginal likelihood of the neutral model ℳ_*n*_, integration is thus replaced by summation over the reference population SFS:

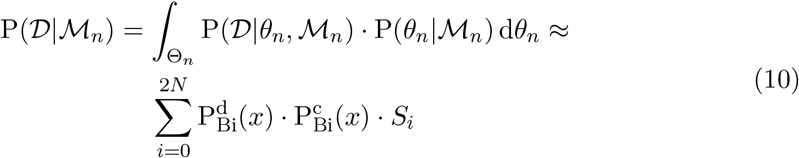

Here, the binomial probabilities are weighted by the fraction *S*_*i*_ ∈ [0, 1] of variants having allele count rank *i* in the normalized reference population SFS. The error-adjusted observation rates *π* entering the binomial terms are computed as in Eq. (5), with the distinction that under neutrality the underlying allele frequency is identical in affected and control cohorts and is given directly by the SFS rank:

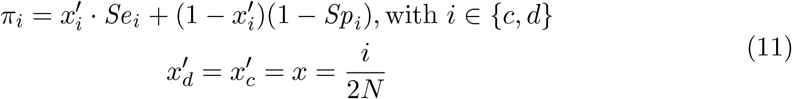

The reference population is characterized by its effective population size *N* ; the same value of *N* is therefore used consistently in both the neutral and pathogenic models, where it enters through Nei’s formulation of the allele frequency distribution.

## 3 Population-specific instantiation of the neutral model

The site frequency spectrum of the reference population for the neutral model cannot be taken directly from existing resources such as gnomAD, as this would introduce circularity and would not provide sufficient resolution to model allele counts across arbitrary sample sizes. Instead, the SFS is generated under a model of neutral mutation and genetic drift, implemented via Wright-Fisher simulations parameterized by the demographic history of the population and a varying mutation rate. Spectra simulated at different mutation rates are then combined by weighting them according to a distribution of *µ*, which may be specified as a continuous density, a discrete set of values, or a fixed constant.

Although discrete mutation rate distributions from previous studies like Seplyarskiy et al. 2023 could in principle be used within this framework, the aim here is to avoid reliance on population-specific external inputs and to demonstrate a self-contained procedure for inferring a contiguous mutation rate model directly from demographic assumptions and high-quality population genomic data. Using gnomAD v4 as an example, this approach shows how the mutation rate distribution and the neutral site frequency spectrum can be instantiated jointly through simulation and data-driven parameter optimization.

The same procedure can, in principle, be applied to other populations as suitable demographic reconstructions and genome-wide allele count data become available, without structural changes to the model.

### 3.1 Simulation of the reference population site frequency spectrum

To simulate neutral site frequency spectra of a large hypothetical human population, a forward Wright-Fisher simulator was implemented in pure R. The simulator assumes complete neutrality and includes only bidirectional recurrent mutation (parameterized by an input value of *µ*), random genetic drift, and prescribed changes in population size. Population history is specified as an arbitrary function of generation number, and the simulation operates directly at the level of per-variant allele counts. The simulator’s source code is available in the project’s GitHub repository.

### 3.2 Mutation rate distribution model

Mutation rate, together with population history, is a primary determinant of the site frequency spectrum observed in genomic data. Demography governs how mutation events are transformed into allele frequencies through genetic drift, whereas heterogeneity in mutation rate across loci determines the rate at which variants enter this process.

When population history is specified, Wright-Fisher simulations define a forward mapping from mutation rate to the expected contribution of variants across allele frequencies. Within this framework, the observed site frequency spectrum can be represented as a mixture of spectra generated at different mutation rates. Although mutation rates at individual loci are not identifiable from frequency data alone, the aggregate mutation rate distribution can be constrained by requiring that this mixture reproduces the observed spectrum under neutrality.

Accordingly, the mutation rate distribution is treated as a latent mixing distribution whose parameters are inferred by optimizing the agreement between simulated and observed neutral site frequency spectra, conditional on a fixed demographic model. This procedure is used to calibrate the allele frequency component of the framework, rather than to infer locus-specific mutation rates.

Mutation rate varies across the human genome by several orders of magnitude and reflects contributions from multiple hierarchical levels, including broad genomic regions, local sequence context, and specific mutational mechanisms (Kong et al. 2012; Aggarwala and Voight 2016; Hodgkinson, Ladoukakis, and Eyre-Walker 2009; Hodgkinson and Eyre-Walker 2011). Despite extensive empirical characterization, no mechanistic genome-wide model currently predicts the full distribution of mutation rates across sites (Hodgkinson and Eyre-Walker 2011). For analytical convenience, mutation rate heterogeneity has been sometimes summarized using simple parametric forms such as a lognormal distribution, with dispersion parameters chosen to reflect observed variability (Hodgkinson, Ladoukakis, and Eyre-Walker 2009; Harpak, Bhaskar, and Pritchard 2016). However, recent base-resolution models that directly incorporate sequence context and molecular determinants of mutagenesis reveal a substantially more complex structure, with at least two well-separated components commonly attributed to baseline mutation processes and highly mutable contexts such as CpG sites (Aggarwala and Voight 2016; Carlson et al. 2018; Seplyarskiy et al. 2023), along with additional substructure near the dominant peak around 10^−8^ (Seplyarskiy et al. 2023).

To capture this bimodality while keeping a number of parameters to a minimum, the mutation rate distribution was modeled as a weighted sum of two lognormal components. This formulation reflects the major empirical features reported in the literature while retaining a small number of interpretable parameters and avoiding overparameterization beyond what can be supported by site frequency data.

### 3.3 Instantiation for gnomAD NFE populations

The proposed approach imposes several requirements on the population dataset used for model instantiation. Population history must be sufficiently well characterized to allow reliable demographic modeling, sample sizes must be large, and total allele numbers must be available not only for variant calls but also for non-called positions, as accurate representation of monomorphic sites (SFS rank 0) is essential. In addition, uniform and consistently high allele numbers across genomic positions are desirable, allowing site frequency spectrum bins to be approximated directly from allele counts without introducing bias from variable coverage; this criterion favors whole-genome data over exomes.

The non-Finnish European (NFE) population satisfies these requirements particularly well, having been extensively studied in terms of demographic history and population structure. In gnomAD v4, the introduction of genome-wide total allele number tables enables accurate construction of the full site frequency spectrum, including monomorphic sites, which was not possible in earlier releases. For these reasons, gnomAD v4 NFE whole-genome data, comprising 34025 samples, were selected as the reference dataset for instantiating the neutral model.

Model instantiation proceeded in two stages. First, the neutral site frequency spectrum expected for the present-day non-Finnish European population was simulated under a forward Wright-Fisher model incorporating mutation and genetic drift, parameterized by a published demographic history. Second, the parameters of the mutation rate distribution were optimized such that binomial samples drawn from the simulated population matched the observed frequency spectrum at neutral sites in gnomAD. The same collection of simulated spectra generated at different mutation rates was then combined, using the optimized mutation rate distribution as weights, to construct the final neutral site frequency spectrum used in subsequent analyses.

To generate simulated site frequency spectra, Wright-Fisher simulations were performed for 250 mutation rate bins spaced evenly on the natural logarithmic scale between 10^−10^ and 10^−5^. Two simulation depths were used for different stages of the instantiation procedure. For each bin, 1 million independent loci were simulated for mutation rate optimization, while 10 million independent loci were simulated to generate the final reference spectra used for downstream modeling, using the European population history as specified in Weghorn et al. 2019. The resulting final effective population size was 4343838 individuals, yielding a symmetric unfolded site frequency spectrum of length approximately 8.6 million.

The target site frequency spectrum was constructed from gnomAD v4 whole-genome dataset. Genome-wide tables of total allele numbers were combined with the corresponding variant site information. Positions were restricted to high-quality synonymous single-nucleotide substitutions using the comprehensive list provided by Seplyarskiy et al. 2023, which enumerates all possible substitution directions passing position-specific quality filters and therefore allows inclusion of monomorphic sites. This set comprised 19969231 candidate variants.

Variants were further filtered to retain those with gnomAD FILTER values of “PASS” or “AC0”, as well as positions absent from the variant call tables, provided that total allele number was at least 99% of the expected maximum (≥ 151668 alleles for the genome dataset). After filtering, 19426609 variants remained. The final target site frequency spectrum was then constructed using NFE allele counts from gnomAD v4 genomes.

Nonlinear optimization was used to estimate the five parameters of the mutation rate model: the mean and standard deviation of each lognormal component and their relative weight in the mixture. Optimization was performed using the 1M-locus simulations described above, while the higher-depth simulations were reserved for construction of the neutral reference spectrum and subsequent likelihood calculations^1^. Trial site frequency spectra were generated from the Wright-Fisher simulations by applying one additional generation of pure genetic drift to subsample the simulated population down to 34025 individuals. Spectra corresponding to different mutation rates were then combined according to the candidate mutation rate distribution and compared to the target gnomAD spectrum using binned summaries.

Comparison was performed using eight site frequency spectrum bins defined over allele counts: [0, 0], [1, 1], [2, 2], [3, 3], [4, 5], [6, 10], [11, 2*N ·* 0.05%] and [2*N ·* 0.05%, 2*N*], with *N* denoting the effective population size of the simulated population. The objective function was defined as the sum of squared differences between simulated and observed bin fractions.

Optimization was carried out using the local gradient-free algorithm BOBYQA (Powell 2009) implemented in the R package nloptr, and was run to convergence. Initialization used a 99:1 sum of two identical lognormal distributions with *µ* = −17, *σ* = 0.5 on log scale, allowing the components to separate if supported by the data.

The resulting mutation rate PDF visibly demonstrates two distinct contributions: a major narrow lognormal component with a peak at *µ* ≈ 7.1 *·* 10^−9^, and a slowly decaying background component which provides non-negligible support for higher mutation rates up to 10^−7^ and above (Fig. 1). This shape resembles previously reported genomewide mutation rate distributions obtained using independent methodological approaches (Aggarwala and Voight 2016; Harpak, Bhaskar, and Pritchard 2016; Kong et al. 2012).

**Figure 1:**
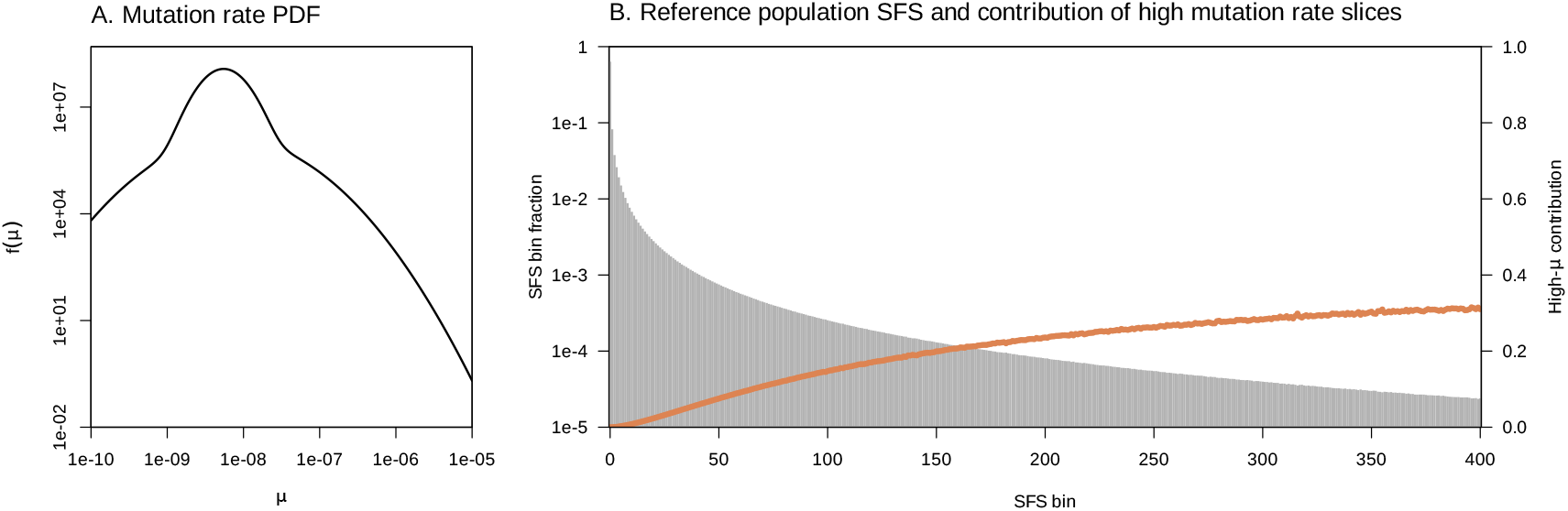
Optimized mutation rate distribution and the corresponding reference population neutral site frequency spectrum. (A) Optimized mutation rate probability density used for discretization into mutation rate slices. Both axes are in log scale. (B) Reference population site frequency spectrum at final effective population size *N* = 4343838, generated under a Wright-Fisher process and weighted by optimized *µ* distribution. The relative contribution of slices generated under high mutation rate (*µ* ≥ 10^−7^) is shown for each SFS bin.

Optimization of *µ* PDF resulted in near-perfect agreement between simulated and observed site frequency spectra at the level of binned summaries in gnomAD v4 NFE genomes (Fig. 2A,B). Agreement was also evident at the level of individual SFS ranks within the low allele-frequency range up to 0.05%, which is most relevant for dominant variant analysis (Fig. 2C,D).

**Figure 2:**
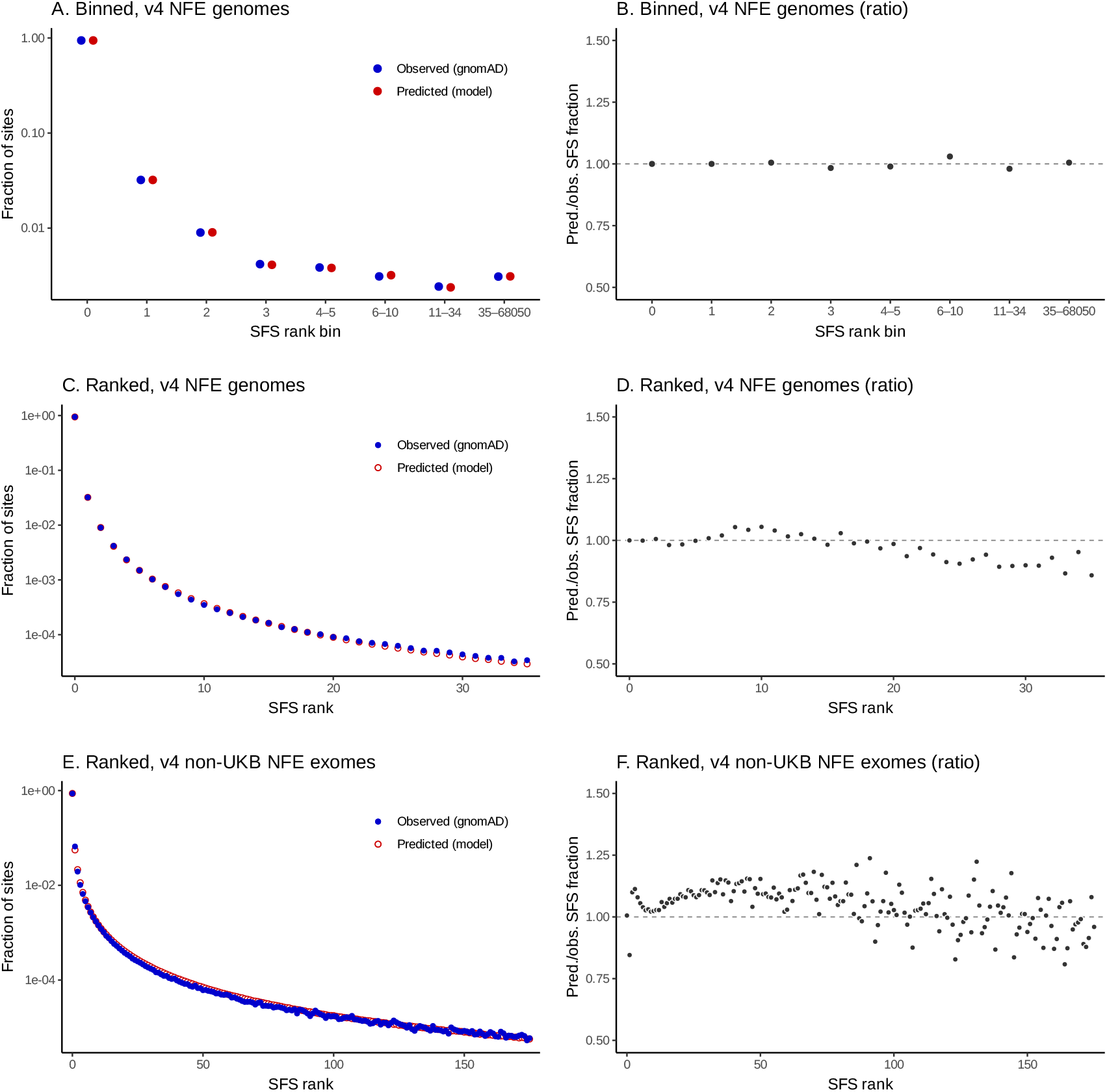
Comparison of observed neutral site SFS fractions with *µ*-weighted simulations in gnomAD populations. Observed site frequency spectra from gnomAD v4 NFE genomes and non-UKB NFE exomes at neutral sites are compared to one-generation drift subsamples of the corresponding sizes derived from WF-simulated reference population under optimized distribution of the mutation rate. Panels (A, C, E) display observed and simulated SFS fractions in log scale; panels (B, D, F) demonstrate predicted-to-observed ratios of SFS fractions. The top row (A – B) presents the binned SFS fractions in v4 NFE genomes used in *µ* optimization. The middle and bottom rows (C – F) present SFS comparisons at the level of individual ranks up to allele frequencies 0.05% for v4 NFE genomes (C – D) and v4 non-UKB NFE exomes (E – F).

Applicability of the resulting reference population model to larger sample sizes was further evaluated by comparing predictions to gnomAD v4 non-UKB NFE exomes (175054 individuals). Using the previously optimized mutation rate distribution and the same set of Wright-Fisher simulation slices, expected spectra for this sample size were generated by an additional generation of pure drift and compared to the observed data. Agreement across SFS bins remained largely preserved within the 0.05% allelefrequency range; however, the singleton class (rank 1) showed a systematic deviation, being underestimated by approximately 15% relative to the observed value. Subsequent low-frequency bins were correspondingly slightly overestimated, consistent with a compensatory effect arising from normalization of bin fractions (Fig. 2E,F).

This excess of observed singletons in the larger sample may reflect limitations of the demographic model, including the absence of migration or population substructure and a possible underestimation of the rate of the most recent phase of rapid population growth. In addition, a fraction of observed singletons may represent residual false-positive calls despite filtering, as the expected allele frequencies of true singletons in samples of this size (∼ 10^−5^ … 10^−6^) approach previously reported sequencing and calling error rate estimates (for details, see Subsection 6.2).

### 3.4 Robustness to deviations from idealized spectra

Comparison to gnomAD v4 non-UKB NFE exomes at a sample size of 175054 individuals revealed a systematic underestimation of the singleton class by approximately 15%, accompanied by a compensatory overestimation of adjacent low-frequency bins. This discrepancy raises a natural concern regarding the use of large external reference datasets as controls: namely, whether deviations of this magnitude in the ultra-rare tail of the neutral site frequency spectrum (SFS) could materially affect the Bayes factor when the same dataset is used to evaluate control allele counts.

To address this question directly, robustness to deviations from the idealized neutral spectrum was assessed using a worst-case perturbation framework directly calibrated to the observed gnomAD discrepancy. Rather than assuming a particular parametric form of misspecification, reference population SFS perturbations were constructed using linear programming (LP) to identify redistributions of SFS mass that minimize or maximize the neutral marginal likelihood, subject to a set of empirically motivated constraints.

Formally, for fixed observed allele counts (*k*_*d*_, *k*_*c*_), the LP objective optimizes the change in the neutral marginal likelihood induced by a perturbation of the reference population SFS, while holding all other components of the model fixed. Only the neutral model is affected by these perturbations; the pathogenic marginal likelihood is unchanged. As a result, worst-case shifts in the Bayes factor arise entirely through changes in the neutral marginal likelihood.

Perturbations were constrained to reproduce the empirically observed differences between predicted and real gnomAD spectra at the non-UKB v4 exome scale (350108 alleles) for the three lowest-frequency bins (ranks 0, 1, and 2), corresponding to invariant sites, singletons, and doubletons. These bins capture both the region where the discrepancy is concentrated and the region to which the marginal likelihood is most sensitive. Total probability mass was conserved, and nonnegativity constraints were enforced to ensure that all perturbed spectra remained valid probability distributions.

To reflect the intended use of gnomAD as a control cohort, the neutral marginal likelihood in the LP objective was evaluated using the same control sample size (*n*_*c*_ = 350108).

To restrict perturbations to a biologically interpretable neighborhood of the baseline spectrum, a per-bin relative deviation bound was imposed, limiting the magnitude of change in each reference population SFS component relative to its nominal value. The primary analysis used a bound of *±*50% per bin, representing a substantial relaxation of the neutral model while excluding degenerate solutions in which entire frequency classes are effectively eliminated. Sensitivity to more permissive bounds was explored separately.

Robustness was evaluated across representative allele count scenarios chosen to probe sensitivity at key interpretive boundaries. These included the absence or presence of a single affected carrier (*k*_*d*_ = 0 or 1), small and large affected cohort sizes (*n*_*d*_ = 2000 and 200000), and models with or without sequencing errors. In all scenarios, a singleton observation in controls (*k*_*c*_ = 1) was considered, as this configuration is maximally sensitive to misspecification of the ultra-rare spectrum. Results are reported as worstcase shifts in log_10_ Bayes factor, derived solely from changes in the neutral marginal likelihood.

Under the *±*50% per-bin bound, the resulting range of Bayes factor changes remains limited in magnitude across all tested scenarios, with maximal shifts in log_10_ *BF* remaining below thresholds corresponding to supporting-level evidence in the ACMG framework (Tavtigian, Harrison, et al. 2020). In several cases the LP solution saturates the imposed constraints, with only negligible additional change observed upon further relaxation of the perturbation bound.

In contrast, in large-cohort, error-free scenarios with a single affected carrier, further relaxation of per-bin bounds permits increasingly adversarial redistributions of SFS mass that satisfy the low-frequency constraints at the gnomAD scale while corresponding to biologically implausible spectra. In this regime, substantially larger shifts in log_10_ *BF* become possible. However, such solutions reflect underconstrained optimization rather than realistic neutral model misspecification and are therefore not considered representative of plausible deviations from the assumed spectrum.

Together, these analyses demonstrate that the Bayes factor is robust to empirically observed deviations from the idealized neutral spectrum at the scale of large control datasets such as gnomAD v4 non-UKB NFE exomes. Sensitivity becomes pronounced only when the admissible perturbation space is expanded to include near-degenerate spectrum configurations well beyond reasonable bounds of biological plausibility.

### 3.5 Reference-anchored site frequency spectrum reconstruction

A site frequency spectrum obtained from gnomAD or similar resources is neither classically folded nor unfolded. Even under neutrality, it is unavoidably *reference-anchored*, because alleles are defined relative to a specific reference genome used for variant calling. In the case of hg38, approximately 72.6% of the reference sequence derives from a single individual (Aganezov et al. 2022), implying that the reference state is not chosen uniformly between the alleles at a polymorphic site.

Under neutrality, at the level of SFS counts rather than individual loci, it is reasonable to approximate the probability of the minor allele being present in a population-derived reference genome as proportional to its population frequency. Under this assumption, reference anchoring can be modeled as a stochastic reassignment of allelic states: for each allele count class in a folded SFS (up to the midpoint), a binomially sampled fraction of variants is reassigned to the symmetric upper class in the corresponding reference-anchored spectrum, with probability determined by allele frequency. This procedure was implemented as a statistical reconstruction of a reference-anchored SFS from a folded spectrum.

As shown in Figure 3, the reconstructed spectrum at the scale of gnomAD v4 non-UKB NFE exome set closely follows the observed SFS in the allele count range where it diverges from the fully folded representation, although some deviations were expected due to partial ancestry mismatch between admixed reference and the population. At very low allele frequencies (*<*1%), the difference between folded and reference-anchored spectra is minimal; nevertheless, this reconstruction step was retained for completeness and was applied to the folded reference population SFS prior to its use in neutral model marginal likelihood calculations, yielding a reference-anchored population spectrum that is compatible with the structure of the observed data.

**Figure 3:**
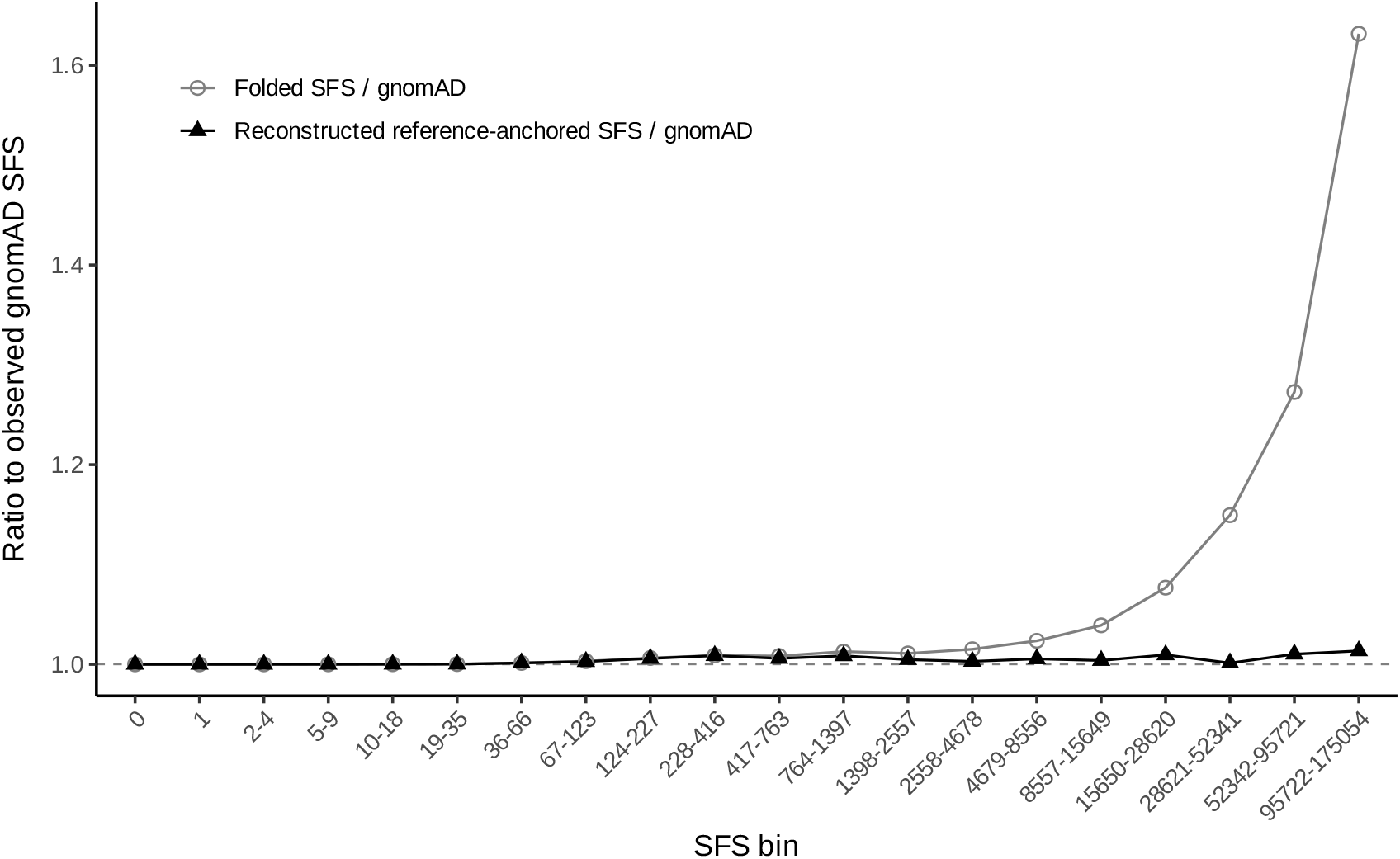
Comparison of folded and reconstructed reference-anchored site frequency spectra relative to the original gnomAD data for synonymous sites. Ratios of binned allele counts in the folded SFS and in the statistically reconstructed reference-anchored SFS to the corresponding binned allele counts observed in gnomAD v4 non-UKB NFE population were computed across logarithmically sized SFS rank bins up to the midpoint of the spectrum. The analysis was performed on a list of putatively neutral synonymous variants as in Subsection 3.3, using the procedure from Subsection 3.5.

## 4 Computational implementation

The Bayesian allele count model was implemented in pure R as a callable function that evaluates the marginal likelihoods of the pathogenic and neutral models and returns their ratio as a log_10_ Bayes factor. The implementation is distributed as open-source software under the MIT license via a public GitHub repository.

At runtime, the Bayes factor function accepts observed allele counts and model hyperparameters, together with a population-genetic formulation of the model. This formulation can be provided in several equivalent forms, reflecting different levels of prior knowledge about mutation rates and site frequency spectra (SFS). In its most general configuration, the function operates on a set of simulated reference population SFS slices generated for discrete mutation rate bins, accompanied by the corresponding bin boundaries. These slices are combined internally according to a user-supplied mutation rate model.

The mutation rate component of the model can be specified in one of three ways. First, a continuous probability density function for the mutation rate may be supplied, together with its hyperparameters; this is the default mode used throughout the present study. Second, a discrete mutation rate distribution may be provided directly as weights over the SFS slices. Third, a fixed mutation rate value may be assumed, in which case the corresponding SFS slice is used directly without averaging. These options allow the same computational framework to accommodate a wide range of modeling assumptions, from fully parametric to simplified or exploratory settings.

For efficiency, when both the mutation rate distribution and the SFS set are fixed, the model also supports the use of a pre-combined SFS representation in which mutation rate averaging has been performed in advance. In this case, the SFS is supplied as a single normalized spectrum, avoiding redundant construction during repeated Bayes factor evaluations.

Numerical integration required for marginal likelihood evaluation is performed using the “hcubature” routine from the R package cubature. The relative error tolerance was set to 10^−10^, no limit was imposed on the number of function evaluations, and the absolute error tolerance was left at its default value.

For reproducibility, the repository accompanying this manuscript includes both the full model implementation and pre-constructed data objects corresponding to the parameterization used in this study, including the set of simulated reference population SFS slices and the optimized mutation rate distribution derived from gnomAD v4 NFE genomes.

### 4.1 Numerical integration and stability considerations

The marginal likelihood of the pathogenic model involves a two-dimensional integral over allele frequency *x* and mutation rate *µ*, together with an additional one-dimensional term arising after analytic singularity cancellation. The two-dimensional integral was evaluated in natural log space over both variables, with the integrand rescaled by the exponential of the summed log-parameters to preserve the original measure. This parameterization substantially improves numerical stability in regimes where the integrand varies over many orders of magnitude or exhibits narrow contribution regions.

For the pathogenic model, the mutation rate integration domain was set to (0, 10^−5^], extending the lower bound relative to the neutral model SFS construction (that used a minimum *µ* slice 10^−10^) while retaining the same upper bound and prior *f* (*µ*). In calculating cubature, the allele frequency domain was taken as (*x*_0_ = 10^−12^, *x*_*m*_ = *R/*(2*K*)], where *R* denotes disease prevalence and *K* penetrance. The upper bound corresponds to the maximal biologically plausible pathogenic allele frequency under the assumption that all affected individuals (and all non-penetrant carriers) are heterozygous. The lower bound is set to a small positive value to limit domain extent while remaining far below frequencies contributing appreciably in the regimes considered.

After singularity cancellation, the remaining one-dimensional term involves integration over *µ* of the mutation rate prior multiplied by a gamma cumulative distribution function evaluated at the upper *x* bound *x*_*m*_. It was computed using adaptive numerical integration with respect to *µ*. A finite *x* bound induces a deficit relative to the corresponding untruncated expression; this deficit enters the final marginal likelihood through the analytic weighting specified in Eq. 8. To ensure that this effect does not materially influence reported results, the induced contribution to the marginal likelihood was monitored at runtime, and execution was halted if it exceeded 1% of the final value.

A diagnostic surface of the two-dimensional integrand in (log *x*, log *µ*) space is generated and exposed in the function call, allowing direct inspection of dominant contribution regions. The current implementation has been empirically verified to be stable across the regimes considered here. Additional numerical safeguards, including inward boundary-shrinkage tests, coarse fixed-grid integration checks in log space, and sensitivity analyses with respect to the lower *x* bound in the two-dimensional term, are planned for future versions but are not required for the analyses reported in this work.

### 4.2 Software implementation and online calculator

In addition to the standalone R implementation, an online calculator was developed to facilitate exploratory use of the proposed model and to allow direct evaluation of Bayes factors under the parameterizations considered in this study. The calculator implements the same Bayesian framework and numerical procedures as described in this manuscript and does not introduce additional model assumptions or approximations.

The online tool uses a fixed population-genetic formulation corresponding to the analyses presented here. Specifically, the neutrality model is based on a precomputed reference population neutral site frequency spectrum, and the mutation rate component is represented by the optimized double-lognormal distribution inferred from gnomAD v4 NFE genomes (see Fig. 1). These components are held constant in the web implementation to ensure consistency with the results reported in this work.

Users may vary observed allele counts in affected and control samples, sample sizes, and key model hyperparameters, including penetrance, disease prevalence, inclusion ratio, selection coefficient, and assay error rates for both affected and control datasets. Default parameter values correspond to those used in the main analyses unless manually changed by the user.

Each evaluation presents the log_10_ Bayes factor together with the marginal likelihoods of the pathogenic and neutral models. In addition, selected intermediate results are provided and visualized, including the diagnostic integrand surface described earlier, to verify that the dominant contribution to the integrand mass lies well within the integration domain and is not driven by boundary effects.

Individual Bayes factor evaluations typically require several seconds of computation. To ensure stable performance under concurrent access, the online calculator incorporates result caching and request queuing. For analyses requiring custom site frequency spectra, alternative mutation rate models, or large numbers of repeated evaluations, the standalone R implementation provided with this manuscript should be used.

The calculator code is available under AGPL-3 license in a separate GitHub repository, and a live instance can be accessed at: https://tools.clinbiolab.org/bayes.

## 5 Results

The analyses below examine how the proposed framework behaves under realistic combinations of allele count data and modeling assumptions. The starting point is an examination of Bayes factor landscapes over affected and control allele counts, which expose the structure of support for the pathogenic and neutral models through their marginal likelihoods. This is followed by an exploration of how observed allele counts are translated into relative support for the competing models under different sampling configurations and error assumptions. The resulting behavior is then compared to existing ACMG allele frequency evidence categories. Finally, the sensitivity of the Bayes factor to individual model parameters is assessed directly, holding allele counts fixed in order to isolate parameter effects.

Unless stated otherwise, all analyses in the Results section use a common reference parameterization representative of a dominant rare disease scenario, with affected and control cohort sizes *n*_*d*_ = 10000 and *n*_*c*_ = 129206, sensitivity of 0.9 in both cohorts, imperfect specificity in controls (1 − 10^−6^) and in the affected (1 − 10^−7^), high penetrance (*K* = 0.95), moderate disease prevalence (*R* = 10^−4^), small but nonzero inclusion ratio (*I* = 0.05), and strong selection against heterozygotes (*h* = 0.95). Parameters varied within individual figures are indicated explicitly, and deviations from this reference set are stated where applicable.

The influence of individual model parameters is examined in detail in subsequent subsections, where their effects on the Bayes factor are explored through targeted sensitivity analyses.

### 5.1 Bayes factor landscapes over affected and control allele counts

Figure 4 illustrates the dependence of the Bayes factor on the observed allele counts in affected cases (*k*_*d*_) and controls (*k*_*c*_) under the common baseline scenario defined above. In addition to the Bayes factor surface, the marginal likelihood landscapes for the pathogenic and neutral models are shown separately, directly representing the structure of support for each model across the (*k*_*d*_, *k*_*c*_) plane.

**Figure 4:**
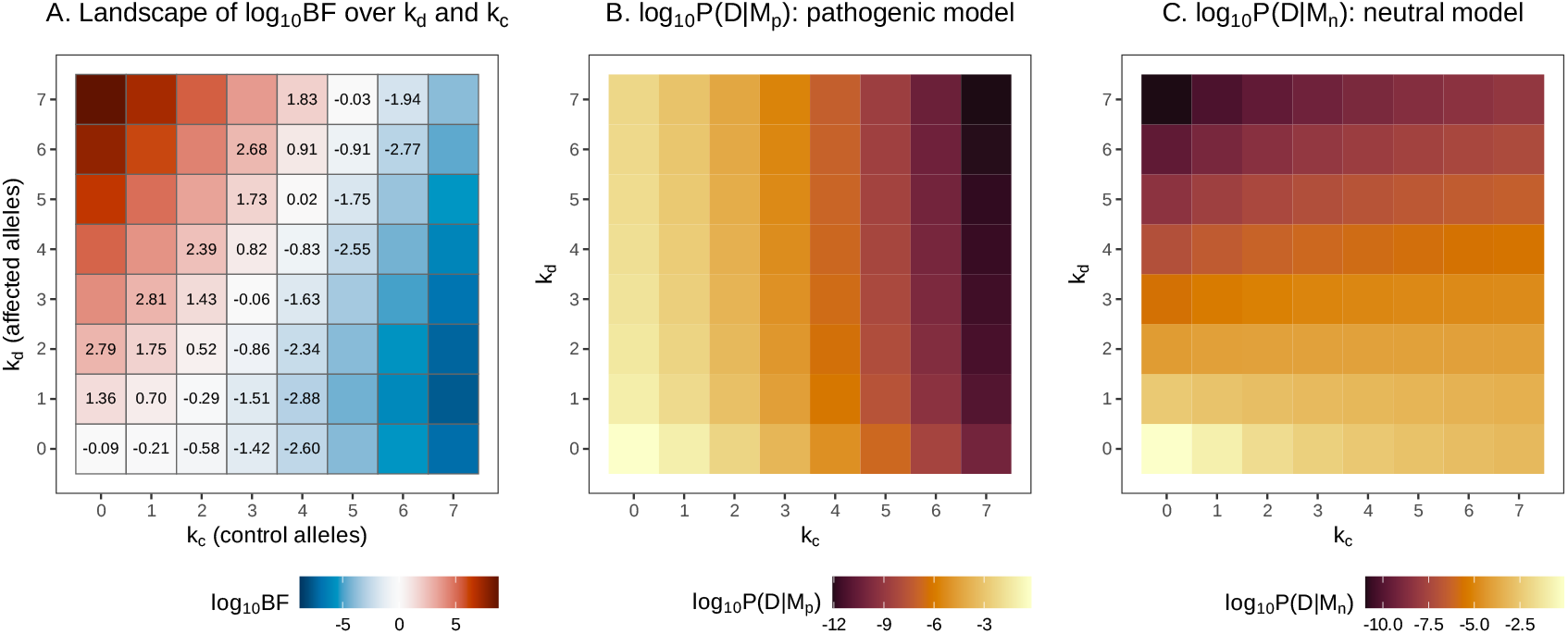
Bayes factor landscape over affected and control allele counts. The dependence of Bayes factor and marginal likelihoods on observed allele counts in affected individuals *k*_*d*_ and controls *k*_*c*_ is demonstrated under a representative dominant rare disease parameterization. Panel A presents the surface of the log_10_ Bayes factor comparing the pathogenic and neutral models across the (*k*_*d*_, *k*_*c*_) plane. Panel B displays the corresponding marginal likelihood surface log_10_ P(𝒟|ℳ_*p*_) under the pathogenic model, and panel C displays log_10_ P(𝒟|ℳ_*n*_) under the neutral model. All panels are evaluated over the same grid of affected and control allele counts using the common default parameters specified in the introduction to the Results section.

Under the pathogenic model, the marginal likelihood decreases monotonically with increasing allele counts in both cohorts (Fig. 4B). Importantly, additional affected alleles do not provide positive support for pathogenicity; instead, they incur a likelihood penalty by shifting posterior mass toward higher mutation rates and higher implied population frequencies. This penalty, however, is comparatively weak for affected counts because the corresponding allele frequency contribution is scaled by disease prevalence and penetrance. In contrast, each additional control allele contributes directly to the inferred population frequency and therefore leads to a steeper reduction in likelihood.

The neutral marginal likelihood surface exhibits a different geometry (Fig. 4C). Rather than being dominated by absolute allele counts, it is structured primarily by the balance between affected and control cohorts, reflecting their exchangeability under neutrality. In the present parameterization, allele counts in the affected cohort exert a stronger influence than those in controls, consistent with the substantially smaller affected sample size and the larger per-allele frequency increment associated with *k*_*d*_.

The resulting Bayes factor landscape (Fig. 4A) reflects the interaction of these two distinct penalty structures. Regions of positive support for pathogenicity are confined to configurations with low control allele counts and moderate affected counts, while increasing *k*_*c*_ produces a rapid and nonlinear shift toward strong support for neutrality. Owing to imperfect control specificity, the initial addition of control alleles at very low counts incurs only a modest penalty, followed by a much steeper decline once the implied population frequency exceeds the range tolerated by the pathogenic model.

Overall, the Bayes factor varies smoothly over the (*k*_*d*_, *k*_*c*_) plane, with no sharp thresholds or discontinuities. The relative impact of affected and control allele counts is parameter-dependent and shaped by cohort sizes, error rates, and disease characteristics, underscoring the inherently graded and context-specific nature of evidence derived from case-control allele observations.

### 5.2 Bayes factor response to observed allele counts under varying scenarios

Figure 5 illustrates how the Bayes factor responds to observed allele counts in affected individuals and controls across a range of cohort sizes and error assumptions. Panels A and C show log_10_ *BF* as a function of the number of affected alleles *k*_*d*_ for several cumulative affected cohort sizes *n*_*d*_, in the absence (*k*_*c*_ = 0) and presence (*k*_*c*_ = 3) of control observations, respectively. In both cases, *BF* increases with increasing *k*_*d*_, indicating progressively stronger relative support for the pathogenic model as affected carriers accumulate. For a fixed *k*_*d*_, larger *n*_*d*_ yields weaker relative evidence, consistent with dilution of case frequency in larger cohorts. Conversely, smaller affected cohorts produce steeper *BF* increases with *k*_*d*_, as the same number of affected carriers implies greater enrichment.

**Figure 5:**
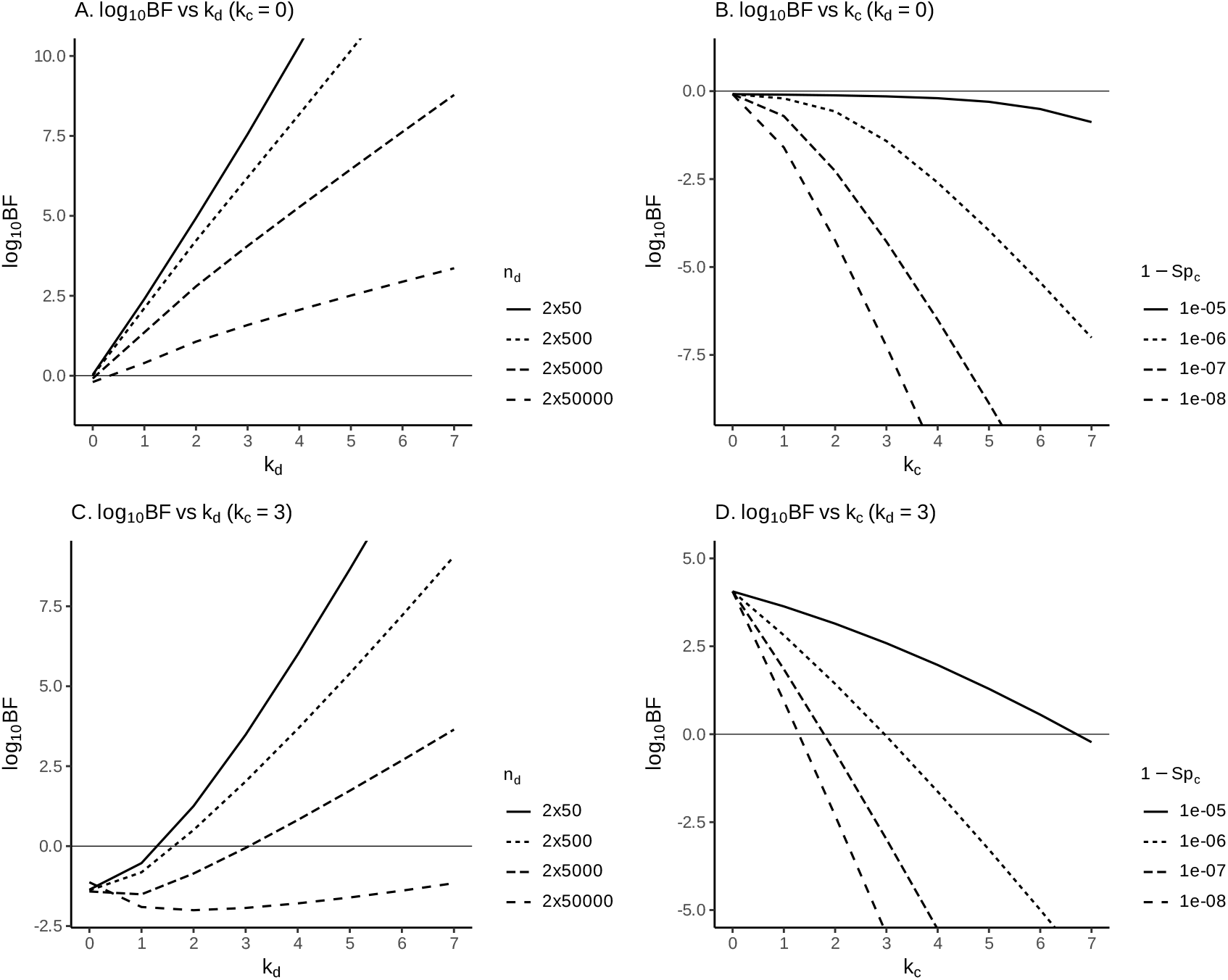
Bayes factor response to observed allele count data. The dependence of the log_10_ Bayes factor on observed allele counts in affected individuals *k*_*d*_ and controls *k*_*c*_ is demonstrated under representative dominant rare disease parameterizations. Panels A and C present log_10_ *BF* as a function of *k*_*d*_ for increasing affected cohort size *n*_*d*_, in the absence (*k*_*c*_ = 0; panel A) and presence (*k*_*c*_ = 3; panel C) of control observations. Panels B and D present log_10_ *BF* as a function of *k*_*c*_ under varying assumptions about false positive rate in controls (1 − *Sp*_*c*_), in the absence (*k*_*d*_ = 0; panel B) and presence (*k*_*d*_ = 3; panel D) of allele observations in the affected. All other model parameters are held at their default values as specified in the introduction to the Results section.

Panels B and D of Fig. 5 show log_10_ *BF* as a function of the number of control alleles *k*_*c*_ under varying control specificity assumptions, again in the absence (*k*_*d*_ = 0) and presence (*k*_*d*_ = 3) of affected carriers. In all cases, increasing *k*_*c*_ decreases *BF*, reflecting loss of relative support for pathogenicity as control observations accumulate. When affected carriers are present (Fig. 5D), this decrease is approximately linear over the plotted range, with the slope determined by control specificity. When no affected carriers are observed (Fig. 5B), log_10_ *BF* remains close to zero for small *k*_*c*_ before declining more rapidly at higher control counts.

Beyond these general trends, Fig. 5C reveals a visible non-monotonic feature when control carriers are present. The mild “dip” observed at very small *k*_*d*_ arises from asymmetric responses of the competing models to rarity. Under neutrality, likelihood is primarily determined by similarity of allele frequencies in cases and controls and does not intrinsically favor zero counts. In contrast, the pathogenic model strongly prefers very low population frequencies through its evolutionary prior, and absence or near-absence of observations in affected individuals is therefore highly compatible with pathogenicity, especially given amplification of population frequency by prevalence and penetrance. As *k*_*d*_ increases from zero to one or two, this intrinsic “rarity advantage” of the pathogenic model is lost before any enrichment signal appears, leading to a transient decrease in *BF* . Importantly, this effect is confined to the negative log_10_ *BF* range and reflects loss of rarity support rather than evidence reversal.

The different behavior of *BF* with respect to control counts at *k*_*d*_ = 0 and *k*_*d*_ *>* 0 arises from the mixture structure of the control call probability in (5): when no affected carriers are observed (Fig. 5B), a small number of control alleles can be accommodated within regions of high likelihood mass where observations are dominated by error terms, whereas in the presence of affected carriers (Fig. 5D) the true-frequency component contributes non-negligibly from the outset, so control counts constrain the underlying population frequency immediately.

### 5.3 Quantitative comparison with ACMG PS4-based frameworks

Table 2 compares the behavior of the proposed Bayesian allele count model with several PS4-based ACMG/AMP implementations (Richards et al. 2015; Bowman 2026; Rowlands et al. 2024; Whiffin et al. 2017) across representative case-control scenarios. All ACMG-style results are expressed as log_10_ odds ratios derived from triggered evidence codes using the Tavtigian point framework (Tavtigian, Harrison, et al. 2020), making them numerically comparable to the log_10_ Bayes factors produced by the present model.

**Table 1:** Worst-case shifts in log_10_ Bayes factor under constrained perturbations of the neutral site frequency spectrum (per-bin bounds *±*50%). Each cell reports [Δ log_10_ *BF* _min_, Δ log_10_ *BF* _max_] for *k*_*c*_ = 1 and *n*_*c*_ = 350108.

| $k_d$ | No sequencing errors | | With sequencing errors* | |
| --- | --- | --- | --- | --- |
| | $n_d = 2000$ | $n_d = 200000$ | $n_d = 2000$ | $n_d = 200000$ |
| 0 | [+0.058, +0.121] | [+0.075, +0.192] | [-0.002, +0.020] | [-0.003, +0.020] |
| 1 | [-0.056, -0.018] | [-0.037, +0.074] | [-0.028, +0.014] | [+0.005, +0.071] |

\* $Se_d = Se_c = 0.9$ , $Sp_d = 1 - 10^{-7}$ , $Sp_c = 1 - 10^{-6}$

**Table 2:**
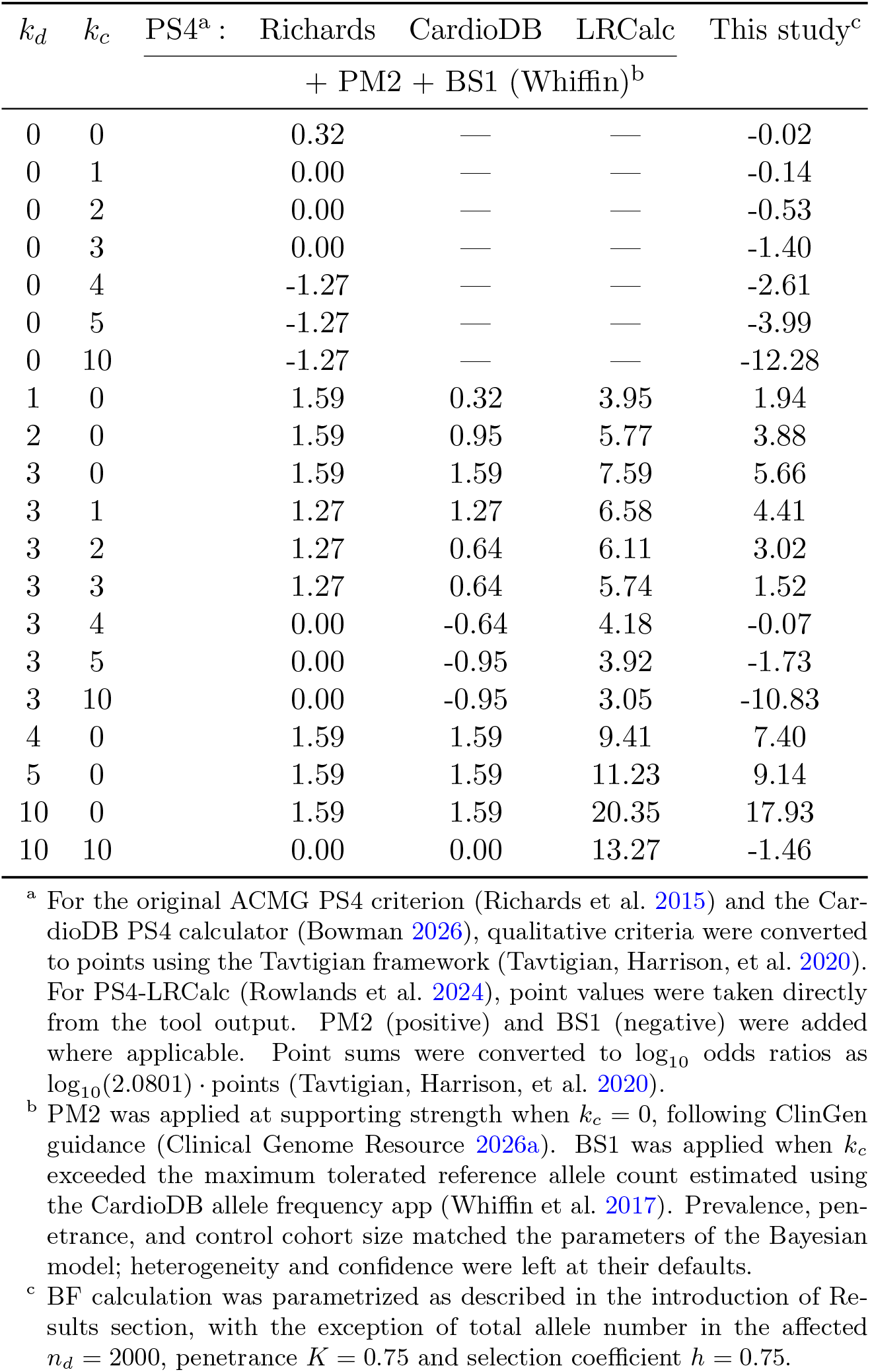
Comparison of Bayesian allele count evidence with ACMG PS4-based frameworks across case-control scenarios. For each combination of affected (*k*_*d*_) and control (*k*_*c*_) allele counts, ACMG-derived evidence strength is reported on a log_10_ odds ratio scale. In the last column, log_10_ Bayes factor is reported.

To ensure that the interaction between case-control enrichment and frequency-based benign evidence is visible within a compact and interpretable set of scenarios, Table 2 was parameterized to yield a non-degenerate range of control allele counts around the maximum credible frequency threshold. In particular, a moderate effective penetrance was used (*K* = 0.75), which places the Whiffin maximum credible allele count at 3 for the chosen control cohort size; higher penetrance values shift this cutoff downward and compress the transition region, reducing the ability to examine behavior on both sides of the threshold. The selection coefficient was constrained accordingly (*h* ≤ *K*) to avoid parameterizations in which selection against carriers exceeds the affected fraction implied by penetrance. Recomputing Table 2 under a higher penetrance baseline (*K* = 0.95, with the corresponding Whiffin cutoff) preserves the qualitative patterns observed here (data not shown).

In the absence of any observed alleles (*k*_*d*_ = *k*_*c*_ = 0), the Bayesian model yields a log_10_ Bayes factor close to zero, reflecting that the data are uninformative with respect to pathogenicity. In contrast, ACMG-based frameworks assign weak pathogenic evidence through PM2 by construction, resulting in a positive log_10_ odds ratio even when no data are available. As control allele counts increase in the absence of cases (*k*_*d*_ = 0), the Bayesian model exhibits a smooth, monotonic shift toward benignity, whereas ACMG implementations transition discretely once the maximum credible frequency threshold is exceeded.

In scenarios with minimal case evidence and no controls (*k*_*d*_ = 1, *k*_*c*_ = 0), the Bayesian model produces evidence strength comparable to the original PS4 criterion, despite no parameters being tuned to reproduce ACMG thresholds. This agreement indicates that the Bayesian framework recovers established ACMG intuition in the lowest informative regime. As additional affected carriers are observed without controls, the Bayesian model accumulates evidence continuously, distinguishing between increasing numbers of affected carriers, whereas PS4-based frameworks rapidly saturate once predefined evidence categories are reached.

The most pronounced differences emerge when a fixed number of affected carriers is accompanied by increasing control allele counts. For example, with three affected carriers (*k*_*d*_ = 3), PS4-based frameworks remain positive or only weakly negative until a frequency threshold is crossed, while PS4-LRCalc continues to assign very strong evidence based primarily on relative enrichment. In contrast, the Bayesian model shows a graded decline in support for pathogenicity and rapidly transitions to negative evidence once control counts exceed levels compatible with the assumed disease prevalence and penetrance. This behavior reflects the tight coupling between case observations, population allele frequency, and control expectations inherent to the Bayesian formulation, rather than reliance on externally imposed frequency cutoffs.

Finally, symmetric scenarios with equal case and control counts at larger scales (*k*_*d*_ = *k*_*c*_ = 10) highlight a fundamental distinction between ratio-based and frequency-constrained approaches. While PS4-based methods converge to neutrality and PS4LRCalc remains strongly positive, the Bayesian model assigns net evidence toward benignity, indicating that absolute allele frequencies, not only relative enrichment, are incompatible with a rare, penetrant disease model.

Overall, these results demonstrate that the Bayesian allele count framework aligns with ACMG PS4 behavior in minimal informative settings, while providing a continuous and biologically constrained quantification of evidence that diverges from threshold-based implementations when absolute allele frequencies become incompatible with rare disease assumptions. The framework thus makes clear how much evidence is present in the data, allowing policy-driven thresholds or categorical interpretations to be applied transparently downstream.

### 5.4 Sensitivity of the Bayes factor to model parameters

Figure 6 summarizes the sensitivity of the Bayes factor to key model parameters under a fixed allele count configuration. Each panel shows the one-way response of log_10_ *BF* to variation in a single parameter, with all remaining parameters held at their default values; default settings are indicated by open circles. All panels share a common *y*-axis range to allow direct comparison of effect magnitudes.

**Figure 6:**
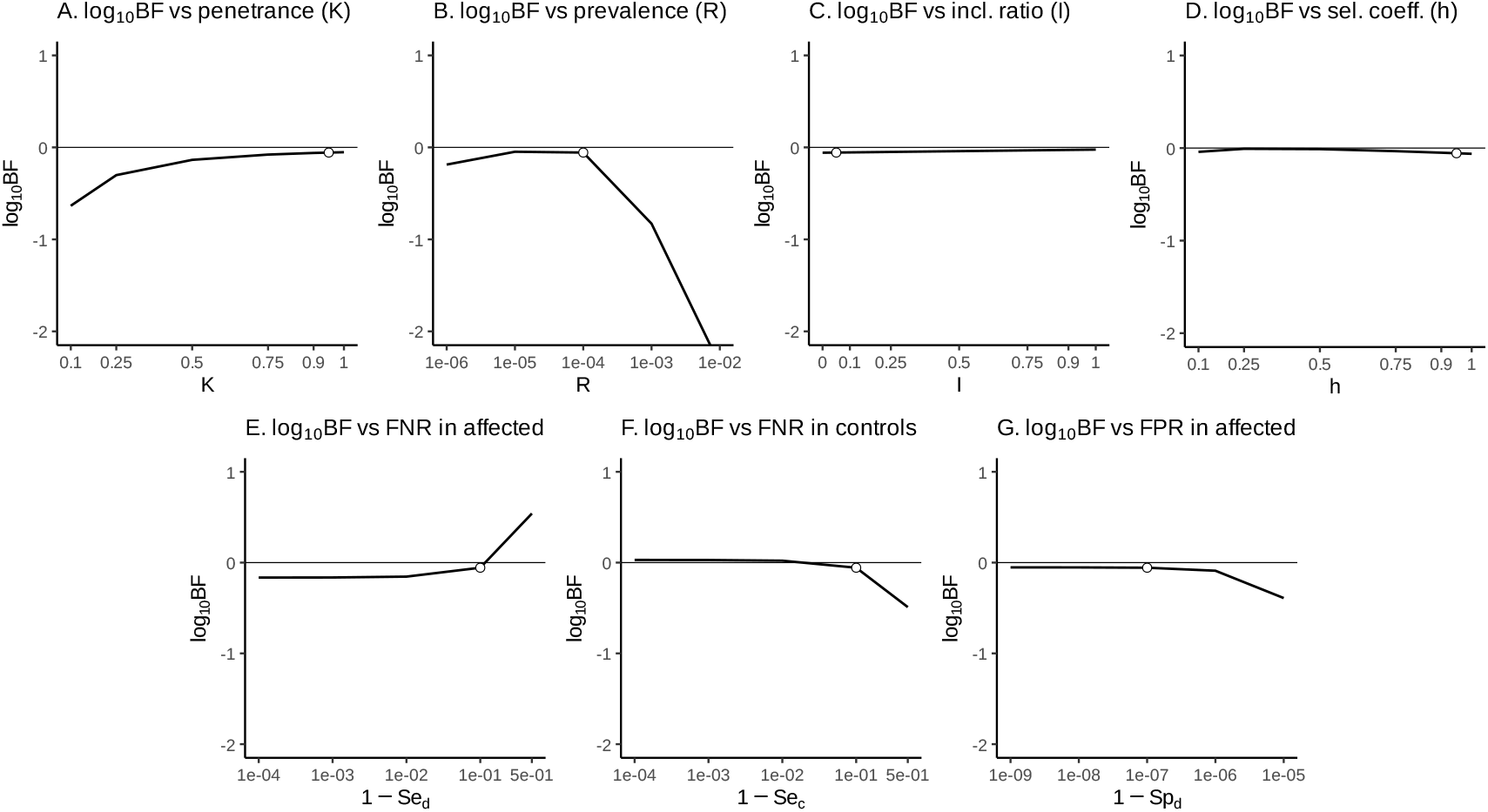
Sensitivity of the Bayes factor to model parameters. Each panel shows the one-way response of log_10_ Bayes factor to variation in a single model parameter, with all other parameters fixed at their default values as specified at the beginning of the Results section. Parameters varied are penetrance *K* (A), disease prevalence *R* (B), inclusion ratio *I* (C), selection coefficient *h* (D), false negative rate in the affected 1 − *Se*_*d*_ (E), false negative rate in controls 1 − *Se*_*c*_ (F), and false positive rate in the affected 1 − *Sp*_*d*_ (G). The default value of the parameter varied in each panel is indicated by an open circle. All panels share a common *y*-axis range to allow direct comparison; curves extending beyond this range are clipped.

The Bayes factor exhibits heterogeneous sensitivity across parameters. Penetrance and disease prevalence have the strongest impact on *BF*, whereas inclusion ratio and selection coefficient produce only minimal changes over the examined ranges. Intermediate effects are observed for assay error parameters in affected and control samples.

Reducing penetrance or increasing disease prevalence consistently decreases the Bayes factor (Fig. 6A-B). In the present borderline regime, lowering penetrance from its default value reduces log_10_ *BF* by approximately 0.6 units, while increasing prevalence leads to substantially larger penalties, exceeding 2 log_10_ units at high prevalence. This behavior follows directly from the structure of the case likelihood (Eq. 6): for a fixed number of observed affected alleles, lower penetrance or higher prevalence shifts the likelihood mass toward larger underlying population allele frequencies. Under the pathogenic model, such upward shifts increasingly strain the mutation-selection prior, whose characteristic scale in the present parameterization is on the order of *µ/h*, leading to reduced model support.

Assay error parameters show asymmetric effects. Increasing the false negative rate in affected individuals (i.e., reducing case sensitivity) increases *BF* (Fig. 6E), because the same observed affected allele count becomes compatible with higher underlying case enrichment under Eqs. (4–6). In contrast, increasing the false positive rate in affected individuals decreases *BF* by introducing spurious case alleles (Fig. 6G). Increasing the false negative rate in controls decreases *BF* (Fig. 6F). The effect of false positive rates in controls is explored separately in Section 5.2.

By comparison, variation in the inclusion ratio and selection coefficient has negligible influence in this allele count configuration (Fig. 6C-D), with absolute changes in log_10_ *BF* remaining below approximately 0.06 across the examined ranges.

## 6 Discussion

In this study, a fully Bayesian quantitative framework was developed for transforming allele count observations into a ratio of support for pathogenic versus neutral models under an autosomal dominant mode of inheritance. The approach formulates the evaluation of allele count evidence as a model comparison problem and allows a wide range of disease characteristics and variant-specific properties to be specified directly as continuous, user-adjustable parameters, rather than being absorbed implicitly into binary criteria definitions. By integrating over uncertainty in allele frequency and mutation rate, the framework produces Bayes factors that directly reflect the relative ability of competing models to explain the observed data.

A defining feature of the Bayes factor in this context is that it is quantitative by construction. Evidence strength is not assigned post hoc via categorical thresholds, but is instead encoded directly in the numerical value of the Bayes factor itself, reflecting the relative support of competing models. This stands in contrast to threshold-based allele count criteria, where continuous observations are ultimately reduced to discrete bins. While any downstream interpretation framework may choose to impose caps, envelopes, or category mappings on Bayes factor values for policy or usability reasons, such constraints are external to the method rather than intrinsic to it. The Bayes factor itself does not require thresholding to be meaningful; it remains a continuous measure of evidence that can be interpreted or transformed as needed without altering its underlying statistical meaning.

For practical instantiation and testing, the method was applied to the non-Finnish European (NFE) population from gnomAD v4. This choice was driven by pragmatic considerations rather than by any assumption of representativeness. The NFE cohort currently provides uniquely large, high-quality public datasets with position-specific allele numbers, alongside a comparatively well-characterized demographic history that can be leveraged for site frequency spectrum modeling despite its complexity and past bottlenecks. At the same time, European populations constitute only a limited subset of global human genetic diversity and cannot be viewed as a proxy for humanity as a whole (Tishkoff et al. 2009). To address this limitation, the framework itself was developed in a population-agnostic manner, outlining a procedure by which it can be instantiated for other populations as sufficiently detailed demographic models and population-scale data become available.

Clinical validation of the proposed framework was deliberately outside the scope of the present work. The intent here is methodological rather than prescriptive: to provide a transparent, explicitly parameterized model that exposes assumptions often left implicit in current practice, rather than to propose an immediately deployable clinical decision rule. Rigorous clinical validation would require coordinated efforts across multiple expert groups, carefully curated datasets, and formal consensus on ground-truth definitions – the conditions that go well beyond what can reasonably be achieved within a single methodological study. Accordingly, the framework is best viewed at this stage as a scientific tool for quantitative reasoning and sensitivity analysis, and as a foundation for future validation efforts rather than as a replacement for existing clinical workflows.

Across a broad range of scenarios explored in the Results section, the qualitative behavior of the Bayes factor is largely consistent with intuitive expectations. However, two parameters that are often treated implicitly or heuristically in allele-frequency-based reasoning emerge as having a particularly strong and sometimes counterintuitive influence on model support: the effective size of the affected cohort, and the rates of assay-level false positive allele observations. The following subsections examine these factors in detail.

### 6.1 Affected cohort size as a dominant but nontrivial determinant of allele count evidence

A noteable result of this work is that the Bayes factor response to observed allele counts in affected individuals depends strongly on the size of the affected cohort. For a fixed number of observed affected alleles, changes in the effective affected sample size can alter the Bayes factor by orders of magnitude. This dependence is therefore not a technical detail but a primary determinant of the strength of allele count evidence.

At first glance, this observation may appear obvious. In many areas of biomedical research, an “affected cohort” implies a deliberately assembled group of patients sharing a narrowly defined disease entity, recruited under uniform inclusion criteria. Rare disease genetics rarely conforms to this paradigm. Monogenic disorders are highly heterogeneous; in addition, the same pathogenic variant may be compatible with multiple phenotypes or exhibit variable expressivity. Moreover, the question addressed by allele count evidence is not whether a variant explains the specific phenotype of an individual patient, but whether the variant can be pathogenic in principle.

For the purposes of pathogenicity assessment, the affected cohort must therefore be defined more abstractly. In this framework, the affected cohort is the set of individuals who (i) have a phenotype compatible with any disease that the variant could plausibly cause if pathogenic, and (ii) were investigated in a manner that would have allowed the variant to be detected and reported had it been present.

Operationally, this definition requires several considerations. First, the phenotypic scope must be defined as broadly as justified by current knowledge of gene-disease associations, encompassing all disorders credibly attributable to pathogenic variation in the gene. Second, the cohort size must reflect not only reported cases, but all cases that *could have been* reported. This includes individuals whose samples were technically suitable for detecting the variant (appropriate sequencing technology, target regions, and variant type), and whose analyses occurred at a time when the gene-disease relationship was sufficiently established for variants to be recognized and communicated. Studies lacking physical access to the variant locus, or predating recognition of the relevant gene-disease association, do not contribute meaningful information and should not be counted.

Importantly, the affected cohort is not restricted to cases in which the variant of interest was identified, or another variant in the same gene. It also includes phenotypically matching cases in which a causative variant in any other gene was found, provided that the variant under evaluation was interrogated and would have been reported had it been present. In addition, it must include phenotypically matching cases that yielded no genetic diagnosis and therefore were never published. Although such negative cases are rarely enumerated systematically, their numbers can often be inferred from known diagnostic yields for the relevant disease context.

Finally, for allele count evidence to be interpreted coherently within a population-genetic framework, the affected cohort must be population-matched both to the control sample and to the population used for instantiating the neutral model. This requirement follows directly from the model structure: both pathogenic and neutral marginal likelihoods are defined with respect to a specific population allele frequency distribution. Mixing affected cases drawn from a different ancestry or demographic background introduces systematic distortion that cannot be corrected by downstream parameter tuning.

Within the proposed Bayesian framework, these complexities do not require explicit modeling of gene heterogeneity, allelic heterogeneity, or phenotype stratification. All such sources of variation are implicitly absorbed into the unknown population allele frequency of the variant, over which the marginal likelihood is integrated. Consequently, when estimating the affected cohort size (*n*_*d*_), any study that genuinely interrogated the variant – regardless of outcome – contributes information. From the perspective of pathogenicity inference, a case harboring the variant, a case solved by another gene, and a negative case constrain the same underlying parameter.

This interpretation turns affected cohort size from a vague reporting concept into a concrete modeling choice. It also clarifies why intuitive definitions of “affected” (for example, equating it with all published cases of a given disease) can distort allele count evidence by simultaneously omitting unreported negative cases and including studies in which the variant under evaluation could not have been reliably detected.

### 6.2 False positive variant calls in population databases

A key determinant of the Bayes factor behavior in the presence of large control cohorts is the assumed false positive rate (FPR) of variant calls in population reference datasets. In the present framework, the control-side FPR enters the observation model as the probability that a non-reference allele is reported for a given chromosome when the true underlying allele at that chromosome is reference. This definition does not require the site to be monomorphic; rather, it reflects per-allele misclassification at the level of individual samples.

Direct empirical estimates of per-allele false positive rates are rare, but a value reported for the ESP6500 project places the per-base error rate at approximately 5.5 *·* 10^−7^ (Fu et al. 2013). Useful order-of-magnitude constraints can be inferred from large benchmarking efforts based on Genome in a Bottle (GIAB) truth sets (Zook et al. 2014). In the PrecisionFDA Truth Challenge V1 (2016), the pipeline achieving the highest precision produced 508 heterozygous false-positive single nucleotide variant (SNV) calls within the benchmark regions (*PrecisionFDA Truth Challenge* 2016). The corresponding high-confidence regions BED file (version 2.19) covered ∼ 2.2 billion base pairs, yielding an average heterozygous false positive rate of approximately 2.3 *·* 10^−7^ per position.

These values should be interpreted cautiously. First, they apply specifically to high-confidence genomic regions and well-behaved single nucleotide variants; indels and variants in low-complexity or low-mappability regions exhibit substantially higher false positive rates (Olson et al. 2022). Second, the cited PrecisionFDA result corresponds to the highest-precision pipeline rather than the best balance of precision and recall. As a result, these values likely represent a lower bound on achievable per-allele false positive rates rather than a typical genome-wide average. Nevertheless, they provide a useful empirical anchor indicating that, for modern pipelines and stringent filters, per-allele false positive probabilities much above 10^−6^ are implausible for typical SNVs.

In modern population resources such as gnomAD variant quality is controlled through joint calling across large cohorts combined with extensive site- and genotype-level filtering, which may further suppress per-site false positive rates for single nucleotide variants in aggregate callsets (Karczewski et al. 2020; Chen et al. 2024). At the same time, largescale databases necessarily operate across heterogeneous genomic contexts, and residual artifacts may persist at specific sites even when the average error rate is low. Any single per-allele error rate estimate should be understood as an average over sites: in practice, some genomic positions will exhibit substantially higher error propensities, while others will be effectively error-free.

In the analyses presented here, the control-side FPR was set to 10^−6^, a deliberately conservative value near the upper end of what is plausibly compatible with modern exome-scale population datasets for filtered single nucleotide variants. This choice avoids implicitly assuming near-perfect specificity in very large control cohorts and allows the qualitative influence of rare artifactual control observations on the Bayes factor to be examined in the low-*k*_*c*_ regime. Lower values, such as 10^−7^ or 10^−8^, reduce the magnitude of this effect but preserve the same qualitative behavior.

The results of this study show that the assumed control-side false positive rate can have a pronounced effect on the resulting Bayes factor once control cohorts reach the scale of contemporary population databases. This sensitivity underscores a general limitation of allele-count-based evidence: assumptions about extremely small error probabilities, which are negligible at modest sample sizes, can materially influence inference once hundreds of thousands of chromosomes are considered. Making these assumptions explicit, and allowing them to be varied, is therefore essential for transparent and interpretable variant evaluation.

Although not exerting the same magnitude of influence on the Bayesian model, incomplete detection sensitivity represents an orthogonal and well-recognized challenge in NGS-based variant discovery. Large-scale clinical data indicate that a nontrivial fraction of known pathogenic variants may be missed even in contemporary diagnostic pipelines (Lincoln et al. 2021). This observation highlights that expectations regarding sensitivity should be treated with similar caution as false positive error rates when specifying model parameters.

### 6.3 General limitations of Bayes factors

An important interpretational nuance becomes most apparent when the Bayes factor is close to unity. A Bayes factor near zero on the log scale indicates that the observed data provide similar marginal support for the pathogenic and neutral models. However, this situation can arise for qualitatively different reasons. In some cases, the data are genuinely uninformative, for example when both affected and control allele counts are zero or extremely small, such that a wide range of underlying allele frequencies remains compatible with both models. In other cases, a near-neutral Bayes factor may result from opposing constraints within the data, such as a nontrivial number of affected observations counterbalanced by control observations that are difficult to reconcile with the same parameter values. Although these scenarios yield similar Bayes factors, they correspond to distinct evidential regimes: absence of information versus internal conflict.

This distinction reflects a general property of Bayes factors and likelihood ratio summaries. By construction, a Bayes factor collapses the data into a single scalar measuring relative support between two specified models, but it does not assess whether either model provides an adequate or internally coherent explanation of the data in an absolute sense. As a result, Bayes factors alone cannot distinguish between data configurations that are broadly typical under both models and those that are atypical under both but penalize the models to a similar extent. In practical variant interpretation, this distinction is often recognized implicitly, as variants with no meaningful evidence and variants with strongly conflicting evidence are both frequently classified as of uncertain significance, despite motivating different analytical responses.

Classical Bayesian methodology provides principled approaches for examining such situations, most notably posterior predictive checks and related model adequacy diagnostics, which assess whether the observed data are typical of the parameter regions that dominate the marginal likelihood under a given model (Box 1980; Gelman, Meng, and Stern 1996). In practical variant interpretation, however, full implementation of such diagnostics is not always necessary, as strongly conflicting observational patterns are directly apparent to expert analysts and are handled through manual review rather than automated criteria. Formal adequacy diagnostics nevertheless represent a natural extension of the present framework for applications that require automated or large-scale analyses.

A more fundamental limitation arises from the binary nature of the Bayes factor itself: it is defined only with respect to a specific pair of competing models. While Bayesian inference can be extended to multi-model comparisons or model averaging, the use of a single Bayes factor necessarily entails selecting one reference model to occupy the denominator. In the present framework, this reference model is chosen to be a fully neutral Wright-Fisher process. This choice does not imply that all variants that are non-pathogenic in the rare-disease sense are evolutionarily neutral; rather, it reflects a deliberate decision to contrast the pathogenic model against the most unambiguous and minimally parameterized baseline available. Accordingly, the resulting Bayes factors quantify support for the pathogenic model relative to this neutral baseline, rather than constituting absolute statements about biological effects or evolutionary fitness.

### 6.4 Conceptual relationship to ACMG/AMP criteria

Given the number of assumptions and limitations highlighted throughout the manuscript, it may appear that practical use of the proposed framework is unlikely to be warranted, even after careful validation. However, it is important to recognize that any fixed system of variant interpretation criteria, including ACMG/AMP v3 (Richards et al. 2015), subsequent ClinGen updates (Clinical Genome Resource 2026a), and gene- or disease-specific modifications by VCEPs (Clinical Genome Resource 2026b), can itself be viewed as a special case of applying an underlying model. The crucial difference is that in criteria-based systems this model is implicit rather than explicit: its parameters are not stated, not adjustable, and often not clearly identifiable.

These implicit parameters do not cease to exist simply because they are not written down. When allele-frequency or case-based criteria are formulated for a “typical disease”, this necessarily assumes typical values for penetrance, disease prevalence, affected cohort size, and the probability of observing affected individuals within control cohorts. The present work was motivated by the goal of replacing such hidden assumptions with a transparent and explicitly parameterized framework that (i) is not a black box, (ii) exposes its underlying assumptions, and (iii) allows these assumptions to be meaningfully explored and adjusted.

The present study formalizes a specific evidential component within variant analysis, rather than addressing the interpretation process as a whole. In general, variant interpretation can be viewed as comprising three conceptually distinct parts: a prior probability of pathogenicity, a mechanism for transforming evidence into posterior support, and a set of decision rules acting on that posterior to determine classification, reporting, and actionability. This work addresses only the second component. Accordingly, the proposed framework is prior-agnostic and does not encode assumptions about baseline pathogenicity rates or prescribe classification thresholds. This separation also enables direct comparison with existing allele count criteria, which have previously been recast as Bayesian odds-modifying components (Tavtigian, Greenblatt, et al. 2018; Tavtigian, Harrison, et al. 2020). Within this context, the correspondence observed under realistic scenario supports the compatibility of the proposed approach with the existing ones, as demonstrated in the Results section 5.3.

While the path to routine clinical use of fully formulated Bayesian models is necessarily gradual, there are already several settings in which the proposed framework can be applied safely and constructively. These include serving as a quantitative second opinion when calibrating existing criteria under alternative biological assumptions, supporting population-genetic analyses of allele frequency evidence, and, more broadly, contributing to ongoing efforts to place variant interpretation on a coherent Bayesian foundation grounded in population genetics and disease biology. Such applications were not feasible at the early stages of clinical variant interpretation but have become attainable with recent advances in large-scale population datasets and modeling techniques.

A common objection to explicitly parameterized models is that key quantities, such as penetrance for ultra-rare diseases or variant-specific error rates, are often uncertain. This concern is valid, but it is not unique to formal models: these quantities are already being assumed implicitly in current criteria-based systems. The advantage of a model-based approach is that only biologically meaningful parameters must be specified, and their influence on inference can be examined directly. When uncertainty remains, parameters can be systematically varied over plausible ranges to assess sensitivity. If conclusions depend critically on poorly constrained regions of parameter space, a conservative interpretation can be adopted by selecting parameter values that reduce evidential strength rather than inflate it.

At the same time, the informal nature of existing guidelines confers an important practical advantage. ACMG/AMP, ClinGen, and related guidelines consistently include caveats and exceptions, acknowledging that no fixed set of rules can accommodate all biological scenarios and directly encouraging expert judgment in complex cases. In contrast, the use of numerical models – even transparent ones – can create a temptation to treat outputs as definitive and to disengage expert oversight. This risk underscores the importance of incorporating mechanisms for manual intervention and error containment when applying quantitative models in practice. Possible safeguards include separating evidence into distinct conceptual blocks with capped contributions, and requiring documented justification whenever different lines of evidence provide strongly contradictory signals. Maintaining such expert-in-the-loop constraints is essential to prevent formal models from being applied beyond their domain of validity.

Taken together, these considerations suggest that formal Bayesian models should not be viewed as replacements for expert judgment, but rather as tools for making existing assumptions visible, testable, and scientifically grounded. When used with appropriate safeguards, they have the potential to improve both the transparency and consistency of variant interpretation without sacrificing the flexibility required in real-world clinical decision-making.

### 6.5 Future directions

The framework presented here is intentionally restricted to autosomal dominant inheritance and to population evidence based on allele counts. While this choice reflects both the current clinical use of such evidence and the desire to keep model assumptions simple and interpretable, several natural extensions emerge.

A primary direction for further development is the formulation of an analogous Bayesian framework for recessive inheritance. In this case, the relationship between allele frequency, selection, and disease manifestation differs fundamentally from the dominant scenario, and the gamma-based approximation derived from Nei’s model is no longer applicable. Modeling recessive variants will require separate treatment of homozygous and compound heterozygous genotypes, as well as a more complex representation of selection against carriers. This extension therefore represents a structural generalization rather than a simple modification of existing parameters.

A related but still nontrivial extension concerns X-linked dominant inheritance. Although some aspects of this scenario are conceptually similar to the autosomal dominant case, sex-specific sampling and hemizygosity introduce additional complexity. In particular, the assumptions underlying simplified dominant selection models do not directly transfer to X-linked loci, requiring special treatment of genotype-specific fitness effects rather than reliance on a single effective selection parameter. As a result, extending the present framework to X-linked dominant variants will require partial reformulation of the pathogenic model rather than a purely mechanical adaptation.

Beyond population allele counts, an important long-term goal is the Bayesian formalization of additional lines of clinical evidence, particularly familial data. Evidence derived from *de novo* occurrence and cosegregation cannot be combined with population-based Bayes factors by simple multiplication, as these evidence types share latent parameters such as the true population allele frequency or mutation rate. A coherent treatment will therefore require joint likelihood models that explicitly account for parameter dependencies. Similar considerations apply to phenotypic concordance between observed cases and gene- or domain-specific disease expectations, although the clinical complexity of phenotype modeling places this well beyond the scope of the present work.

The proposed framework also enables systematic evaluation of existing allele count criteria used by Variant Curation Expert Panels. By introducing clear assumptions about penetrance, disease prevalence, and error rates, it becomes possible to assess whether published thresholds correspond to consistent levels of evidential support across different disease contexts. Such analyses may help identify regimes in which current criteria are overly conservative or permissive, and provide a quantitative basis for future calibration efforts.

Finally, the neutral model underlying the Bayes factor calculation can be instantiated for additional populations as demographic histories become better characterized and large-scale population datasets continue to grow. Importantly, the modular structure of the framework allows not only population-specific site frequency spectra but also alternative mutation rate models to be incorporated in a coupled manner. Because mutation rate heterogeneity and population demography jointly shape the site frequency spectrum, a mutation rate model inferred under one demographic assumption cannot be assumed to remain compatible when paired with a different demographic scenario. Future extensions will therefore require empirical validation (and, where necessary, joint calibration) of mutation rate models together with demographic scenarios against the target dataset, rather than direct reuse of components inferred in unrelated contexts.

These directions highlight that the present work should be regarded as a step toward a fully quantitative, Bayesian treatment of variant interpretation, rather than as a complete solution. By directly modeling the evidential structure of allele counts, the framework provides a basis upon which additional inheritance modes, evidence types, and population contexts can be integrated in a principled and transparent manner.

## Data Availability

The R implementation of the Bayesian allele count model, including the Wright-Fisher simulator and population-specific NFE data objects, is available at https://github.com/clinbiolab/allelecount-bayes. The source code for the web-based calculator can be found at https://github.com/clinbiolab/allelecount-bayes-ui.

## Acknowledgements

I am grateful to all my colleagues at ICBL for their patience, kindness and support during the preparation of this manuscript.

AI-assisted tools were used during manuscript preparation for language editing and stylistic refinement. The author reviewed and edited the final text and takes full responsibility for all scientific content, analyses, and conclusions presented in this work.

## Footnotes

1 Repeating the mutation rate optimization using 10 million loci per mutation rate bin produces parameterizations with similar objective values and residual patterns, but with a substantially narrower low-*µ* component peak. When the neutral marginal likelihood is evaluated by discrete summation over a fixed *µ* grid, a narrow shape concentrates weight on a small number of *µ* bins, reducing robustness to *µ* grid discretization. For this reason, the smoother 1M-based parameterization was retained as the default mutation rate prior for all analyses.

